# Efficient small-world and scale-free functional brain networks at rest using k-nearest neighbors thresholding

**DOI:** 10.1101/628453

**Authors:** Caroline Garcia Forlim, Siavash Haghiri, Sandra Düzel, Simone Kühn

**Affiliations:** University Medical Center Hamburg – Eppendorf, Clinic and Policlinic for Psychiatry and Psychotherapy, Martinistraße 52, 20246, Hamburg, Germany; University of Tübingen, Department of Computer Science, Sand 14,72076, Tübingen, Germany; Max Planck Institute for Human Development, Lise Meitner Group for Environmental Neuroscience, Lentzeallee 94, 14195 Berlin, Germany

**Author notes:** indicates a shared first-authorship (Caroline Garcia Forlim, Siavash Haghiri).

**Keywords:** brain network, resting-state fMRI, functional connectivity, k-nearest neighbor, small-world networks, scale-free networks

## Abstract

In recent years, there has been a massive effort to analyze the topological properties of brain networks. Yet, one of the challenging questions in the field is how to construct brain networks based on the connectivity values derived from neuroimaging methods. From a theoretical point of view, it is plausible that the brain would have evolved to minimize energetic costs of information processing, and therefore, maximizes efficiency as well as to redirect its function in an adaptive fashion, that is, resilience. A brain network with such features, when characterized using graph analysis, would present small-world and scale-free properties.

In this paper, we focused on how the brain network is constructed by introducing and testing an alternative method: k-nearest neighbor (kNN). In addition, we compared the kNN method with one of the most common methods in neuroscience: namely the density threshold. We performed our analyses on functional connectivity matrices derived from resting state fMRI of a big imaging cohort (N=434) of young and older healthy participants. The topology of networks was characterized by the graph measures degree, characteristic path length, clustering coefficient and small world. In addition, we verified whether kNN produces scale-free networks. We showed that networks built by kNN presented advantages over traditional thresholding methods, namely greater values for small-world (linked to efficiency of networks) than those derived by means of density thresholds and moreover, it presented also scale-free properties (linked to the resilience of networks), where density threshold did not. A brain network with such properties would have advantages in terms of efficiency, rapid adaptive reconfiguration and resilience, features of brain networks that are relevant for plasticity and cognition as well as neurological diseases as stroke and dementia.

**Highlights:**

- A novel thresholding method for brain networks based on k-nearest neighbors (kNN)
- kNN applied on resting state fMRI from a big cohort of healthy subjects BASE-II
- kNN built networks present greater small world properties than density threshold
- kNN built networks present scale-free properties whereas density threshold did not

## 1. Introduction

Our brain can be thought as a complex network. This view is supported by histological, structural and functional studies, showing complex levels of interaction in the brain. Complex networks applied to neuroscience, hypothesize a network created from inter-connected brain areas, brain areas are called nodes and their connections are described as links (Fornito, Zalesky, and Bullmore 2016; Bullmore and Sporns 2009). Local interconnections can form clusters. Clusters play an important role as functional segregators, processing specialized information and keeping this information at a certain location (Rubinov and Sporns 2010; Bullmore and Sporns 2009). Nevertheless, these centers of specialized information processing are most likely not isolated from each other, thus, one expects intercommunication among the clusters, so that, the information can exchanged in the brain. The connections involved in global intercommunication among different functionally segregated regions would play the role of so-called functional integrators. Moreover, it is of advantage that all process described above are preserved in cases of external or internal perturbations.

An interesting scientific question is, what an optimal brain network would look like or phrased differently: What would be the most efficient way that the brain could work? The concept of small-world organization in networks was introduced by Watts and Strogatz (1998), when they investigated biological, technological and social networks (Watts and Strogatz 1998). This concept is very broad and appears in many other fields and disciplines where complex networks are used, e.g. the power grid networks, social networks, computer science, biology (Telesford et al. 2011; Amaral et al. 2000). It has been suggested that the optimal brain network is topologically reflected by having high clustering and short paths connecting the clusters among themselves (Bassett and Bullmore 2006). This points to the fact that it is equally important to have centers processing specialized information, as it is to have fast communication between them.

To keep these processes properly working in face of external or internal perturbation, the brain network should be resilient, that is maintaining its activity in the face of faults. Thus, besides the small-worldness characteristics, it is desirable that brain networks show scale-free properties. A network is scale-free if its degree distribution follows a power-law function. This property has been associated with network resilience (Cohen et al. 2000; Albert, Jeong, and Barabási 2000; Albert and Barabási 2002). There are multiple studies on brain networks which suggest the presence of scale-free networks (Eguíluz et al. 2005; van den Heuvel et al. 2008). A brain network with such characteristics would bring advantages as both segregate and integrated information processing and enables rapid adaptive reconfiguration, which is important for plasticity and cognition (Baronchelli et al. 2013). Besides, it is very plausible that the brain would evolve to minimize energetic costs of information processing, that is, maximize efficiency (Hilgetag and Goulas 2016; Bassett and Bullmore 2006).

Brain networks can be extracted using various neuroimaging measures: correlations of both functional (e.g. functional magnetic resonance imaging (fMRI), electroencephalography (EEG)) and structural data (e.g. diffusion tensor imaging (DTI), structural MRI). The focus of this article is on networks based on functional correlations from resting state fMRI (rsfMRI). In rsfMRI, subjects lie in the scanner at rest without performing a specific task so that the spontaneous activity of the brain can be measured. Brain regions are said to be functionally connected and therefore, exchanging information, if they present temporal correlation.

Typically, connectivity between areas or voxels is inferred using Pearson’s correlation coefficient, resulting in a large connectivity matrix, sizing number of brain areas/voxels * brain areas/voxels. The connectivity can serve as a distance function between brain regions, where high connectivity values are interpreted as short distance between pairs of nodes and vice-versa, low connectivity values are interpreted as long distances. Nevertheless, there is no consensus on how to treat these connectivity values, resulting in a major problem in brain network analysis: the network construction.

A common practice in network construction is to set a threshold to define where nodes are connected and where they are not. The most frequently used thresholds are called absolute and density threshold, where respectively, only connections that surpass a fixed given connectivity strength are kept or stronger connections matching a given density are kept. Once the full connectivity matrix is thresholded, the final network can be weighted or binary. Weighted networks are those where the connectivity strength is kept, whereas binary ones, the connectivity strength is set to 1. Binary networks are often used in neuroscience and therefore we focus our study in them. In addition the graph measurements, needed to compute the small-worldness, are also defined solely for the binary networks. Noteworthy is that, no thresholding method is unbiased, and there is no consensus about the optimal way to choose a certain threshold. Usually an educated guess is recommended, based on the network features one aims to stress (van Wijk, Stam, and Daffertshofer 2010). The most frequently applied threshold in network neuroscience is the density threshold because it ensures that the networks are constructed with the same budget (the same wiring cost). However, networks emerging from this construction method do not always present the characteristic of being optimally efficient and resilient, namely the small-world and scale-free properties.

Here, we propose an alternative network construction method borrowed from machine learning, called k-nearest neighbor (kNN) graph construction (Luxburg 2007; Daitch, Kelner, and Spielman 2009). Numerous methods in machine learning are based on graphs. The fundamental issue of graph construction also emerges naturally: there is a set of items and a distance function between the pairs of items. The question is how to connect the items by edges and form a graph, in such a way that the resulting graph is informative for machine learning tasks such as clustering, classification, etc. This problem has been extensively explored in machine learning and various solutions were developed (Ng, Jordan, and Weiss 2002; Zhou et al. 2004; Daitch, Kelner, and Spielman 2009; Luxburg 2007). When using kNN to construct a network, the main difference from the density or absolute threshold is that the selection of links is based on local structure and similarities, that is, links that belong to the same neighborhood. In the kNN method, each node of the network is allowed to have exactly *k* connections to its nearest neighbours. In other words, for each node, the *k* strongest connections are kept. kNN had been previously used in neuroscience in algorithms for classification (Arbabshirani et al. 2013; Suárez Sánchez et al. 2014; Zhu et al. 2007), MRI segmentation of the brain (Gang et al. 2013), EEG-based assessment of neurophysiological changes (Kortelainen, Väyrynen, and Seppänen 2011) and in combination with minimum spanning tree graph construction (Alexander-Bloch et al. 2010).

In this paper, we formally introduce the kNN graph construction method for the construction of brain networks. Then, we show the advantages of this method. For that, we applied kNN to a big cohort of healthy participants (N=434) of the Berlin Aging Study II (BASE-II) subjected to resting state fMRI. We show that the kNN network construction leads to the optimally efficient network with small-world characteristics. Moreover, we show that these networks are scale-free, whereas the networks resulting from the traditional network construction do not show this property.

## 2. Methods

### 2.1. Participants

MR eligible participants were recruited within the Berlin Aging Study II (BASE-II, https://www.base2.mpg.de/en; for additional information about the cohort see (Bertram et al. 2014), resulting 438 healthy adults (20-80 years old). None of the participants was on medication that may have affected memory function or had a history of head injuries, medical (e.g., heart attack), neurological (e.g., epilepsy), or psychiatric disorders (e.g., depression). Four subjects were excluded due to excessive movement calculating using framewise displacement (FD) (FD>0.5 (Power et al. 2012)) during the MRI acquisition, resulting in a total of 434 healthy participants.

### 2.2. Data acquisition

Brain images were collected on a Siemens Tim Trio 3T scanner (Erlangen, Germany) using a 12-channel head coil. A T1-weighted magnetization prepared gradient-echo sequence (MPRAGE) based on the ADNI protocol (TR=2500ms; TE=4.77ms; TI=1100ms, acquisition matrix=256×256×176; flip angle = 7°; 1×1×1mm^3^ voxel size) was used to obtain structural images. Whole brain functional resting state images were collected over a period of 10 minutes by using a T2*-weighted EPI sequence sensitive to BOLD contrast (TR=2000ms, TE=30ms, image matrix=64×64, FOV=216mm, flip angle=80°, slice thickness=3.0mm, distance factor=20%, voxel size 3×3×3mm^3^, 36 axial slices). Participants were instructed to look at a fixation cross and relax during data acquisition.

### 2.3. Resting state fMRI preprocessing

To ensure for steady-state longitudinal magnetization, the first 5 images were discarded. First, the acquired data was corrected for slice timing and then realigned, followed by corregistration between structural individual T1 images and functional images. Segmentation into gray matter, white matter, and cerebrospinal fluid were performed. Data was then spatially normalized to stereotactic space of the Montreal Neurological Institute (MNI) and spatially smoothed with a 6-mm full width at half maximum [FWHM]) to improve signal-to-noise ratio. Afterwards, motion and signals from white matter and cerebrospinal fluid were regressed. Data was filtered (0.01 – 1 Hz) to reduce physiological high-frequency respiratory and cardiac noise and low-frequency drift and, finally, detrended. All steps of data preprocessing were performed in MATLAB 2012b (www.mathworks.com) using SPM12 (Wellcome Trust Centre for Neuroimaging, London, United Kingdom) except filtering that was applied using the REST toolbox (Song et al. 2011). Additionally, to control for the effect of motion on functional connectivity measures, the voxel-specific mean framewise displacement were calculated (FD; Power and colleagues (Power et al. 2012). Subjects with FD higher than the recommended threshold of 0.5 were excluded (N=4).

### 2.4. Functional connectivity matrix

To build the resting state functional connectivity matrices, nodes and connectivity strength must be defined. Here the nodes were created based on the AAL atlas (Tzourio-Mazoyer et al. 2002), cerebellum was excluded, resulting in 90 regions of interest and therefore 90 nodes. The node-averaged time series were extracted for each subject using the REST toolbox (Song et al. 2011). As explained above, the BOLD signal was corrected for motion, white matter and cerebrospinal fluid, as nuisance signals of no interest. Connectivity strength was obtained using Pearson’s correlation coefficient resulting in a 90 × 90 matrix. Connectivity values that were not statistically significant (p-value > = 0.05) were excluded.

### 2.5. Graph construction

The goal was to construct an informative binary graph from the connectivity matrix. Here we applied the kNN method to functional connectivity matrices constructed as described in 2.4. In addition, we compared graphs built with kNN to those built based on a density threshold, the most common method in neuroscience.

The density threshold method considers a particular threshold for each subject in such a way that only the highest connections are kept until the desired density is reached. In turn, the kNN method connects each node to its k-nearest neighbors. The k-nearest neighbors of a node are *k* nodes with the highest connectivity values to the node. This procedure can lead to non-symmetric graphs. When denoting the set of k-nearest nodes to node *i* by *KNN*(*i*), if *j* ∈ *KNN*(*i*), it does not necessary mean that *i* ∈ *KNN*(*j*). However, the connectivity matrix is symmetric, and it is desirable to have a symmetric graph as well. To solve this issue, we connected a pair of nodes *i* and *j*, if *i* ∈ *KNN*(*j*) or *j* ∈ *KNN*(*i*). Note that the edges in the final graph have no weight and the final graphs in both density threshold and kNN methods are binary graphs.

In order to obtain a fair comparison, we built density threshold and kNN networks with the same number of edges, considering that the number of edges is the budget for graph construction. In other words, we assumed that the budget of the network to make connections (edges) was fixed and identical for both methods. The value of the threshold controls the number of connections in the density threshold graphs. The constant *k* plays the same role for the kNN graphs. By increasing *k*, we let each node have more connections and the total number of edges in the graph will increase. We chose the range of *k* and the density threshold, in a way that resulting graphs had the same number of edges.

Increasing *k* only by one drastically increases the number of edges in the graph, therefore we cannot fine-tune the number of edges in the kNN method. On the other hand, by slightly changing the threshold value of the density threshold graphs, we can obtain the desired number of edges. Therefore, we chose a fixed range for the parameter k of the kNN graphs. For each value of the parameter *k* and the resulting kNN graph we computed the total number of edges that the graph uses. Then, we tuned the threshold such that density threshold graphs also had the same number of edges. As the number of nodes was fixed, the average degree of graphs was the same across the two methods.

In order to thoroughly understand the behavior of graph constructions we checked a wide range of the parameter *k* ∈ {6,7, …, 60} equivalent to density thresholds from 0.14 to 0.80. In this way, we investigate a range from very sparse to almost fully connected graphs.

### 2.6. Graph Theory analysis

Please note that both graph construction methods in section 2.5 produce binary (unweighted) graphs and the further analysis of small-world and scale-free property is defined only for binary graphs.

#### Graph density

Density of a graph with *n* nodes and *E* edges is defined as

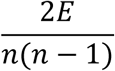

#### Degree

Degree of a node is defined as the number of connected edges to the node.

#### Clustering coefficient

Clustering coefficient of a node was first introduced by Watts and Strogatz (Watts and Strogatz 1998). The clustering coefficient of the node *i* is defined as the number of triangles that it makes with its neighbors divided by the maximum possible triangles. If node *i* has *k* neighbors, then it can form at most *k*(*k* − 1)/2 triangles with its neighbors. Let *Δ* (*i*) denotes the number of triangles that node *i* makes with its neighbors, then the following shows the formal definition of the clustering coefficient:

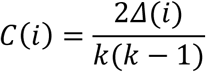

The average clustering coefficient of a graph is defined as the arithmetic mean of the clustering coefficients of nodes in the graph.

#### Characteristic path length

If *d*(*i, j*) denotes the length of the shortest path between nodes *i* and *j*, then the characteristic path length of a graph is defined as the average of distance between all pairs of nodes, formally:

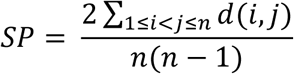

Where *n* denotes the number of nodes in the graph. If two nodes are not connected, then we set their shortest path length to the maximum shortest path length between connected pairs of nodes in the graph.

### 2.7 Small-world network evaluation

The small-worldness of a graph is defined using the average clustering coefficient and the characteristic path length of the graph divided by the equivalent measures in a random graph of the same density. The comparison to the random graph is defined by the following two parameters:

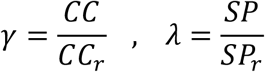

Where, *CC* and *SP* respectively denote the average clustering coefficient and the characteristic path length of the network and *CC*_*r*_ and *SP*_*r*_ are the same quantities for the random graph with the same density. The small-worldness is defined as the following ratio:

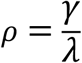

Initially, Watts and Strogatz proposed to use Erdős–Rényi random graphs as a base random model (Watts and Strogatz 1998). In the neuroscience literature, however, it is suggested to compare random graphs with the same degree distribution to have a fair definition of small-worldness (Sporns and Zwi 2004; Maslov et al. 2002). Therefore, we constructed 10 random surrogate networks with the same degree distribution as the initial network. The parameters *CC*_*r*_ and *SP*_*r*_ are the average over the set of 10 random graphs. Similar to other small-world studies in neuroscience (Fornito, Zalesky, and Bullmore 2016; van den Heuvel et al. 2008; Eguíluz et al. 2005), each random surrogate network was built by the following procedure:

We started with the initial network and performed a random rewiring step on edges which preserves the degree of nodes. Next, we chose a set of 4 nodes at random {*i, j, k, l*}, such that the edges *i* → *j* and *k* → *l* exist but the edges *i* → *l* and *k* → *j* do not exist. Note that if the network is very sparse or densely connected, it is not possible to find a quadruple of nodes with the above property. Since we had networks out of these two extreme cases, we could always find a proper set of four nodes. Then we removed the two existing edges *i* → *j* and *k* → *l* and formed two new edges *i* → *l* and *k* → *j*. In this way the connections are randomized but the degree of involved nodes is not changed. The rewiring was done 1000 times with random quadruple of nodes. Considering the number of nodes (90), this number of rewiring steps was enough to produce a random graph.

### 2.8 Scale-free network evaluation

If *p*(*d*) represents the empirical probability of having a node with degree *d*, then the degree distribution of a scale free network should follow a power-law, meaning that *p*(*d*) = *βd*^−*α*^. In this formulation, α is a constant parameter which we refer to as the exponent, and β is a constant to make sure the function represents a probability mass function. In scale-free networks the exponent range should be 2<α<3 (Girvan et al. 2007; Choromański, Matuszak, and Miękisz 2013).

The typical approach to investigate the presence of power-law degree distribution is through the log-log plot representation of the degree distribution. A linear log-log plot indicates a power-law distribution. The R-squared of the linear regression can represent how much of the variance can be expressed by a linear model. If the R-squared is close to one, we can conclude a power-law degree distribution, and thus the presence of scale-free network. Moreover, the slope of the log-log plot represents the exponent parameter *α*, which needs to be in the desirable range (2<α<3) for scale-free networks.

One can obtain the log-log degree distribution plot for each subject and calculate the R-squared value and the slope of the regression curve. However, we intended to also illustrate the average distributions over subjects. Therefore, we presented the results of the scale-free evaluation in two parts. First, we illustrated the average (over subjects) results, then we reported the estimated values of parameters for the whole sample.

Here we explain the procedure to compute the average degree distribution over the subjects: Let assume we estimate the parameters α and β by a linear regressor, for each subject. We denote the estimation of parameters for the subject *i* by 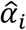 and 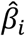, which are respectively the slope and the intercept of the estimated linear regressor for the log-log degree distribution plot of subject *i*. The two parameters can vary significantly among subjects. Some studies compute the average distribution over all subjects in order to estimate a final α (van den Heuvel, Stam, Boersma, & Hulshoff Pol, 2008). In an arithmetic mean over the degree distributions, the subjects who have higher β values have more contribution to the final mean. In this way the final estimation of α can be biased. Here we propose a slightly different approach to overcome this issue. We do not compute the average over distributions, but instead, we compute the average over logarithm value of distributions. Note that by taking the average over log *p*_*i*_ (*d*) values, the final plot will represent: 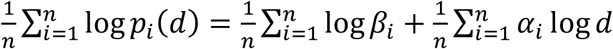. The slope of this plot is the average over the slope values of subjects, 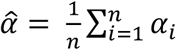.

## 3. Results

In this section, we reported the results of two graph construction methods: density threshold and kNN. The comparison between the two construction methods was done based on two criteria. The first criterion was small-worldness, as efficient networks show small-world properties (Hilgetag and Goulas 2016; Bassett and Bullmore 2006). Secondly, we checked whether the two graph construction methods can produce scale-free networks which is related to resilience of the network (Cohen et al. 2000; Albert, Jeong, and Barabási 2000; Albert and Barabási 2002).

### 3.1. Small-world comparison

First, we reported the two parameters, clustering coefficient ratio γ and characteristic path length ratio λ (Fig 1) for both graph construction methods. The average value of the parameters was calculated over all subjects and the standard deviation was depicted by means of error bars. Note that in each vertical point of the plot the parameter *k* and the density, chosen for the two methods, lead to graphs with the same number of edges, hence the comparison between the two graph construction methods can be considered fair. Moreover, γ and λ were calculated using random graph with the same density.

**Fig. 1.**
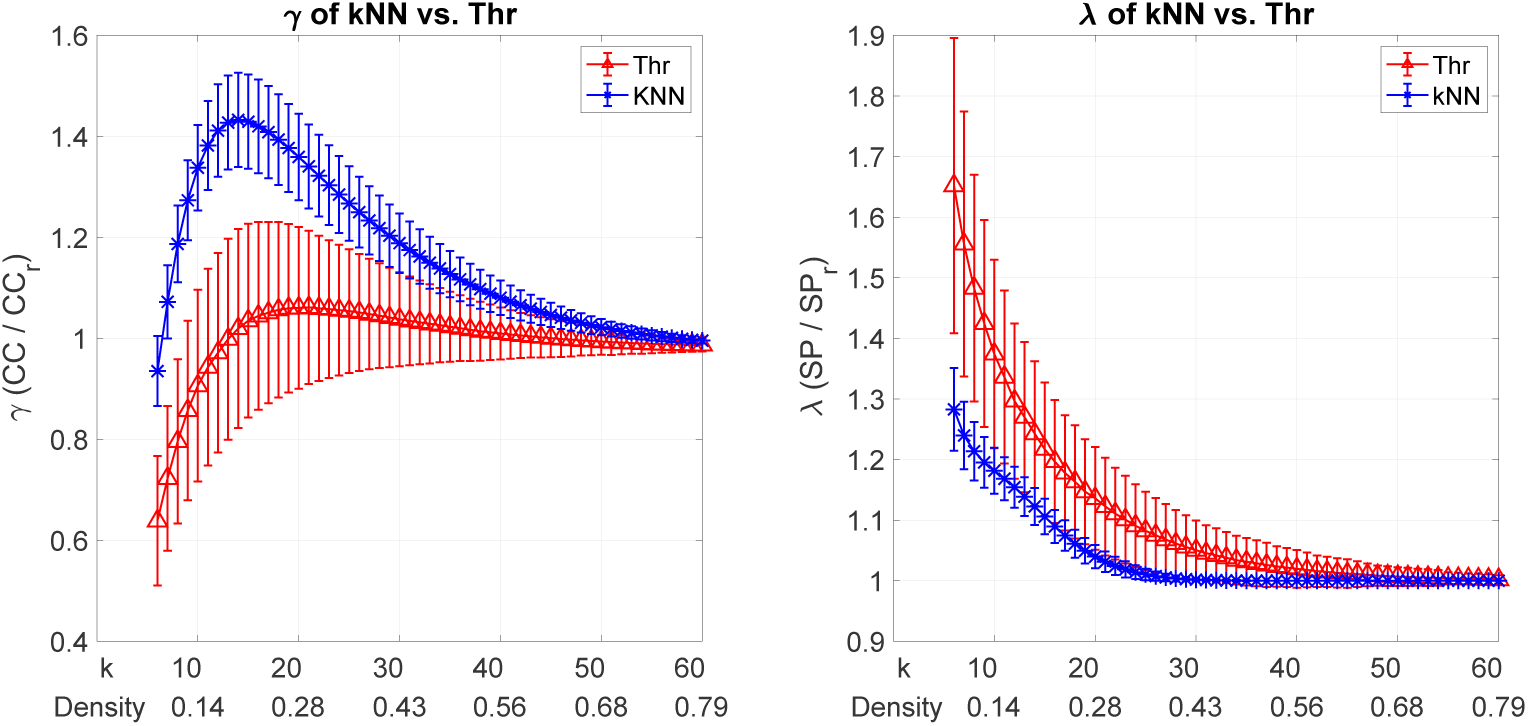
Comparison of parameters (*γ*) and (*λ*) on the left and right respectively, for two graph construction methods: density threshold (Thr in red) and the kNN (in blue). The x-axis of the plots denotes the parameter *k* and density for the kNN and density threshold methods respectively. The errorbars show the standard deviation over subjects. The clustering coefficient ratio (*γ*) is greater for the kNN graph, while the characteristic path ratio (*λ*) is smaller.

The clustering coefficient ratio γ (Fig 1, left) was significantly higher than 1 for the kNN-built graphs (in blue) across all thresholds, meaning more clusters than those for random graphs, especially for *k* values between 8 and 30. While the density threshold graphs presented clustering coefficient ratio close to 1 and below those for the kNN method. This suggests having more well-clustered graphs with the kNN method as compared to density threshold. The characteristic path length ratio λ (Fig 1, right) for the kNN-built graphs showed longer path lengths than random graphs for *k* < 23, however it showed nearly equal characteristic path lengths to the random graphs for *k* > 23. Density graphs produced longer characteristic path length than kNN-built ones. Both parameters suggest that the kNN method will produce networks which have higher small-worldness.

As next step, we calculated the small-worldness (*ρ*) (Fig 2). The kNN-built graphs showed on average higher small-worldness than density threshold ones, especially for *k* < 30. The most efficient networks were constructed with *k* ≈ 18, showing the highest *ρ*. Moreover, the error-bars indicate less variability in the kNN method, that is, a consistent behavior of the kNN method over the set of subjects.

**Fig. 2.**
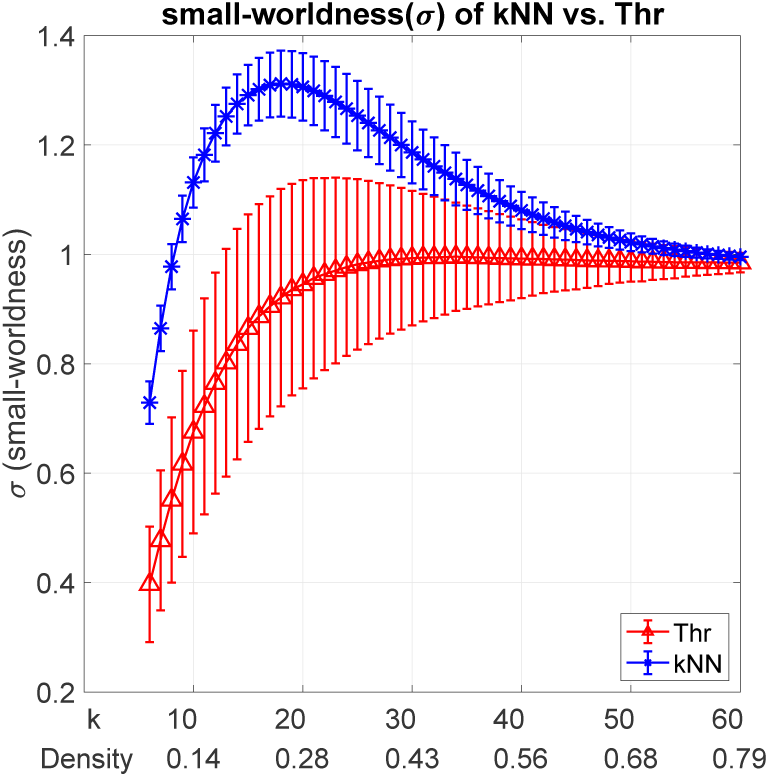
The comparison of small-worldness (***ρ***) for two graph construction methods, density threshold (in red) and the kNN (in blue). The parameters ***k*** and density are shown in the x-axis of the plots. The error-bars denote the standard deviation of the small-worldness with different subjects. The kNN graphs showed significantly higher small-worldness, in particular for ≈ **18**, in comparison to density-thresholded graphs.

### 3.2. Scale-free comparison

We plotted the average (over subjects) logarithm of probability mass functions versus the logarithm of degree, for *k* values by increasing in steps of 10 (Fig. 3). The top row shows log-log plots of the kNN methods, while the bottom row depicts the same plots for the density threshold.

**Fig. 3.**
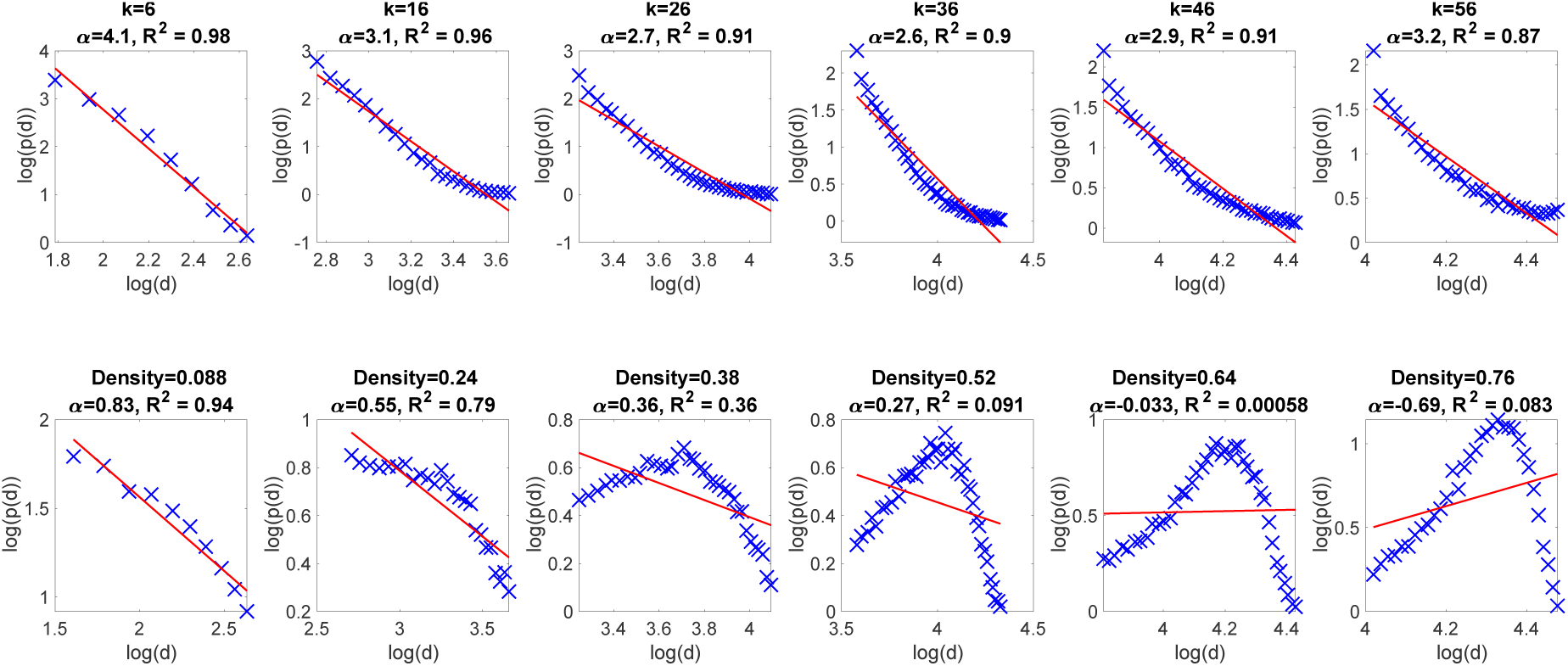
The log-log plots of the average distributions over the subjects. The x-axis is the logarithm of the degree, while the y-axis is the logarithm of the empirical probability of having a node with this degree. The first row shows the results of kNN graph construction method, while the second row shows the results of density threshold graphs. The parameter ***k*** and the density are denoted at the top of each plot. In each column, the graphs have the same number of edges. The R-squared value of the regression and the estimated slope ***α*** is also shown at the top of each plot. The kNN graphs (top row) fit to a linear estimation, moreover, they have exponents (***α***) in the range of scale-free networks. The density threshold graphs are not linear for most thresholds.

In all log-log plots for the kNN method a linear fit was very close to the actual points. Moreover, the R-squared values (denoted at the top of each plot) were close to one. This result suggests that the degree distribution is a power-law *p*(*d*) = *βd*^−*α*^. In addition, except for very sparse graphs (*k* = 6), the whole range of kNN graphs had estimated exponents in the range of 2 < *α* < 3 that characterizes scale-free networks. In comparison, the density threshold method did not show a linear behavior in the log-log plots (Fig. 3-bottom row), particularly with increasing threshold values. Moreover, the estimated exponent *α* failed to fall within the desirable range of scale-free networks.

We also reported the summary statistics of the scale-free test for the whole sample. We computed the R-squared of the linear regression for each subject with the fixed graph construction parameters *k* and density. Moreover, the exponent *α* was estimated based on a linear regression. Fig. 4 (top row) shows the population of R-squared values for the kNN and density threshold methods. We showed see that the kNN method, especially in case of sparse graphs (*k* < 20) presented high R-squared values, which suggests a proper linear fit to the plot. Conversely, the R-squared values were mostly lower than 0.5 for the density threshold method, which means that a linear regressor cannot express the function depicted in the log-log plot. The bottom row of Fig. 4 depicts the population of exponent *α* for the two methods. Again, for the kNN method most of the subjects had exponents in the desirable range while the density threshold method failed to do so.

**Fig. 4.**
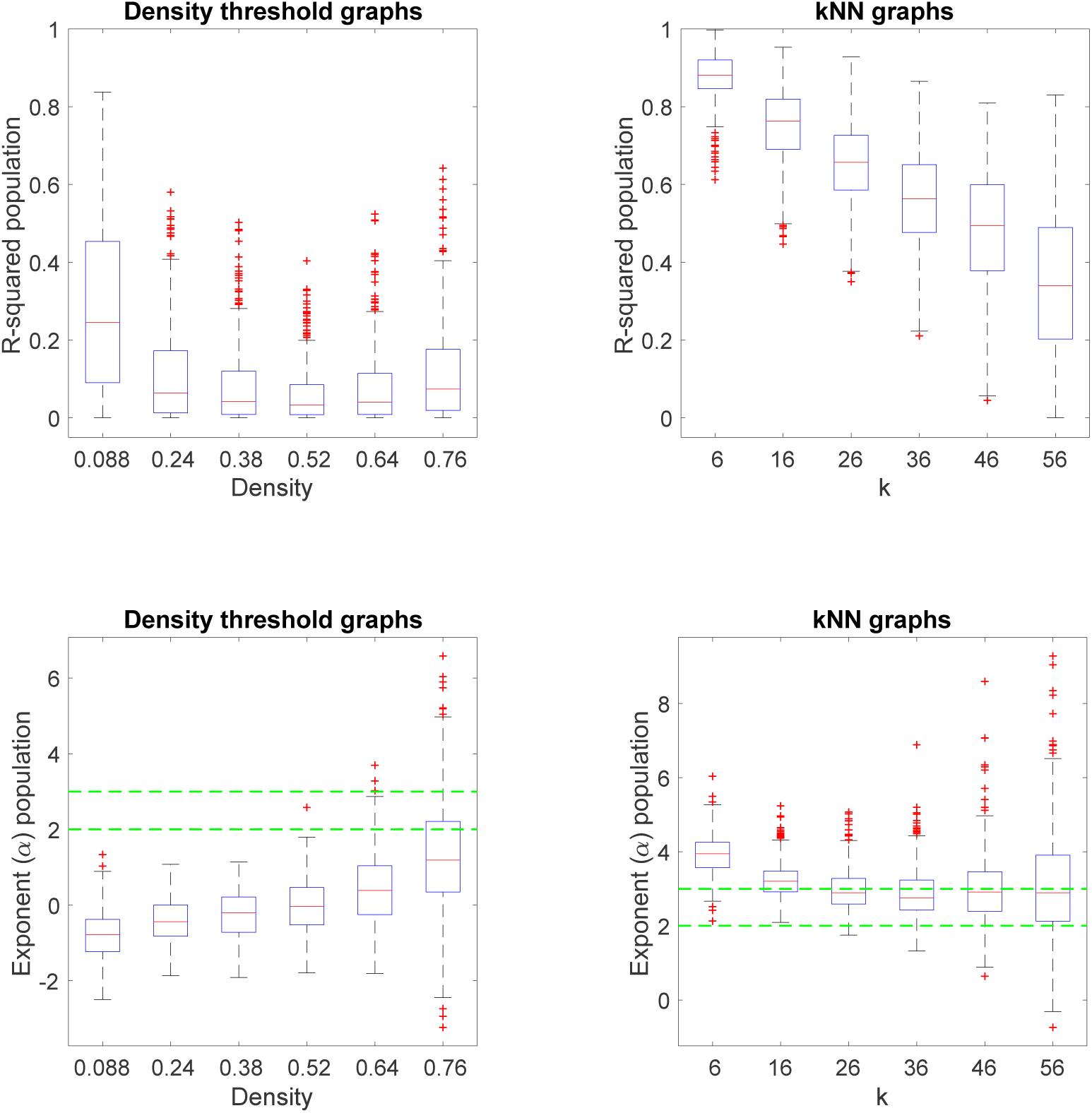
The box-plot of the regression output values, R-squared and exponent ***α***. Each box represents the population of R-squared or the estimated parameter ***α*** for the regression of log-log plot. The central mark in the box is the median, the edges of the box are the 25th and 75th percentiles, while whiskers goes to the maximum and minimum values. Outliers are depicted as red plus signs. The R-squared values are significantly higher for the kNN graphs, suggesting a better linear fit. In addition, the exponent (***α***) is in the range of scale-free network for kNN graphs, with ***k*** > **20**, while this is never the case for density threshold method.

### 3.3. Comparison of networks constructed using KNN and density

Here we present an illustration of the two graph construction methods by plotting the final brain network of a single subject when KNN is applied in contrast to density threshold method (Fig. 5). The parameter k, was chosen to be *k* = 16 for the KNN graph. The corresponding density threshold, for the density threshold graphs, was chosen such that both methods produced the same number of edges (same wiring budget). The choice of *k* = 16 was based on the highest small-worldness (Fig. 2).

**Fig. 5.**
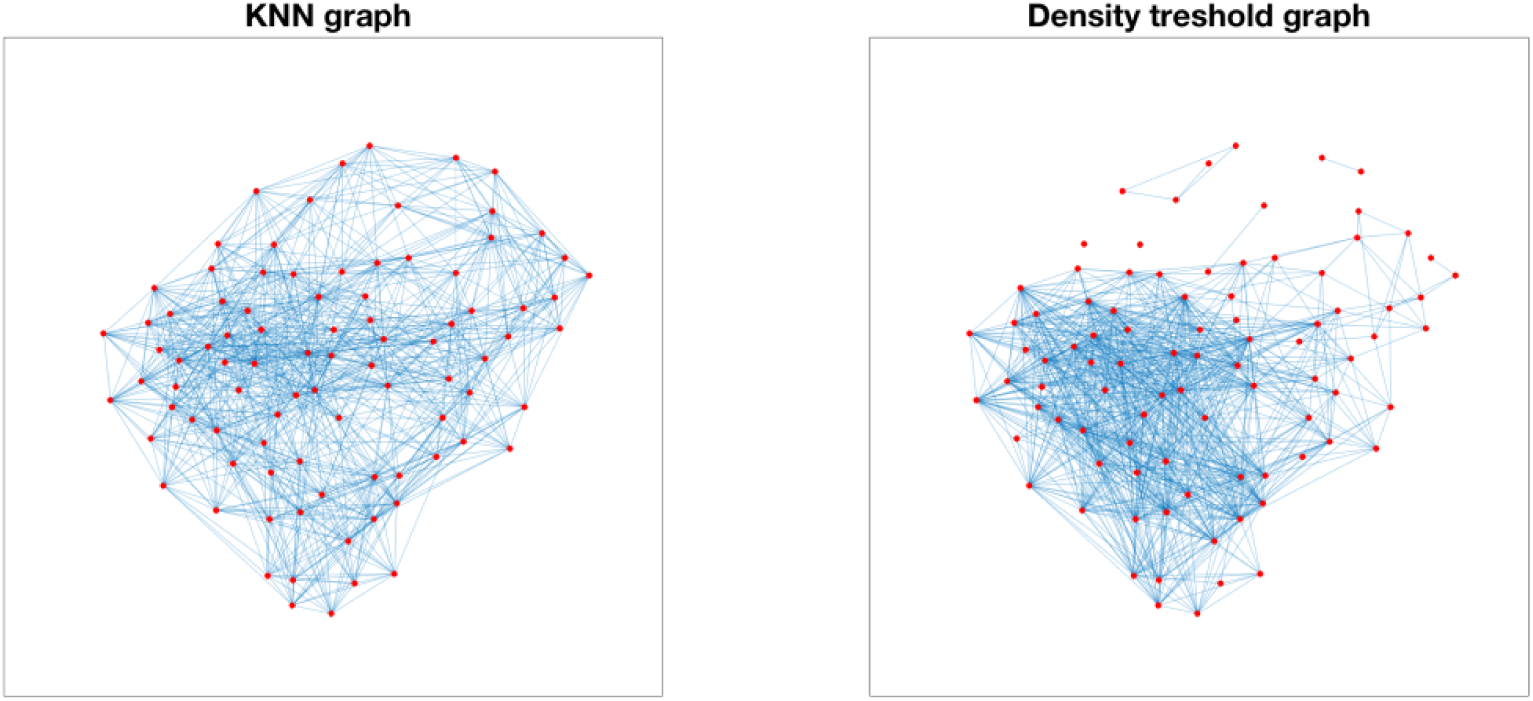
The KNN (left) and density threshold (right) graphs for a single subject.

The main difference observed in the two network construction methods was related to the total interconnectivity of the brain areas (nodes). The graph built using the density threshold method (Fig. 5 – right) is not connected, thus nodes without any edge connection were observed. On the other hand, the KNN graph (Fig. 5 – left) is connected.

The same interconnectivity pattern were seen for other subjects and for the whole range of k parameter values. The reason for such pattern relies on the fact that the density threshold graph is intrinsically prone to produce unconnected components. In brief, the density threshold method uses a global threshold to make edges. In this way, if some of the nodes have systematically weaker connections they cannot form any edge. In other words, if the scale of connectivity values differs among the different regions of the brain (nodes), the density threshold method gives no opportunity to some areas of the brain that have weaker connection, to form an edge. A non-connected graph might not be a favorable structure for the human brain, as some areas of the brain would be isolated and therefore not taken part in the global information processing. In contrast, the KNN method ensures that every node is connected to *k* neighbors. This condition leads to connected graphs for values of *k* which are not extremely small. For a detailed discussion please see (Luxburg 2007).

## 4. Discussion

Our brain is a complex organ with different levels of interaction and connections, from local cellular action potentials to information travelling across the body in our nervous system. The brain network is formed by interconnected areas called nodes and their connections are known as links. Due to the nature of the interactions in the brain, the optimal network should be able to process locally specialized information as well as to connect localized information centers between one another, so that information can travel globally and fast in the brain. A network with such features is said to present small-world characteristics (Watts and Strogatz 1998). Small-world networks work with maximal efficiency and therewith low energetic cost (Bassett and Bullmore 2006). These optimal networks are reflected topologically by having high clustering and short paths connecting the clusters. Beside the efficient organization of networks based on small-world properties, the resilience of the brain network is also desirable. A resilient network is less vulnerable to random perturbation or loss of connections. Such networks exhibit power-law degree distribution and are called scale-free networks.

In practice, brain networks can be inferred using multiple neuroimaging measures such as fMRI, EEG, fiber tracts in DTI, etc. Here, we focus on rsfMRI. In rsfMRI we infer a functional brain network at rest, in which, nodes are voxels or anatomical brain areas and links are the connectivity between the nodes, resulting in a fully connected matrix. In network analysis, connectivity is interpreted as a distance function between nodes: higher connectivity represents short distances, that is, a closer relationship implies strong connectivity. Nevertheless, there is no consensus on how to treat these connectivity values: a fully connected graph may contain spurious connections, e.g. due to noise in data acquisition (Fornito, Zalesky, and Bullmore 2016). However, which connections should be included? This question leads to a major problem in brain network analysis, the network construction.

It is common practice in neuroscience to construct a network, by using global thresholds, such as absolute and density thresholds, the latter being the most popular. When applying a density threshold, the number of links (and consequently the density) is fixed and the stronger connections are kept until the desired density is achieved and the remaining links are set to zero. Considering groups of subjects, the number of connections is the same across subjects which is an advantage when comparing topological properties between groups, however, the final network after thresholding will depend on average strength, for example, subjects with lower average fully connected matrix will have low-weight connectivity links, whereas, for subjects with high average fully connected matrix, only high-weight links will be included. Therefore, when applying global thresholds where nodes are treated the same, as the case for the density threshold, if the connectivity values of specific brain areas are systematically weaker than those for other areas, global thresholds give no opportunity to that particular nodes to make connections in the final graph.

Thresholding in brain network construction is addressed in detail in (van Wijk, Stam, and Daffertshofer 2010). As a final consideration, there is no consensus on network construction methods as there is no unbiased thresholding. Therefore the thresholding method should be chosen carefully by the researcher and should reflect the network features one desires to stress (van Wijk, Stam, and Daffertshofer 2010).

Here we proposed an alternative thresholding method, called kNN (Luxburg 2007; Daitch, Kelner, and Spielman 2009). kNN is used in machine learning, where the problem of graph construction also emerges. kNN connects items that belong to the same neighborhood by allowing the k nearest neighbors of each node. The remaining links, that are not neighbors, are set to 0. In neuroimaging, the connectivity strength can be also thought as the distance function between brain areas. In rsfMRI more specifically, it represents a functional distance and, therefore, is related to the exchange of information in the brain: regions exchanging information have high connectivity values. In turn, spurious connections would present lower connectivity since there is no real relationship between regions. Thus, brain areas can assume the role of the items that must be grouped based on proximity defined by exchange of information.

In this paper, we applied kNN and density thresholding methods to functional connectivity matrices from rsfMRI of a big cohort of healthy subjects. We showed that networks built using kNN method presented small-world and scale-free properties, that is efficient and resilient networks. Our interpretation is that kNN method can preserve key connections and yet reduce the average shortest path length. Shortest paths are essential to the efficient functionality of any network promoting fast communication among specialized centers of information processing.

Regarding efficiency and resilience in the brain, it has been argued that it is plausible that the brain would evolve to minimize energetic costs of information processing (Hilgetag and Goulas 2016; Bassett and Bullmore 2006). In addition, a brain working in small-world scale-free regime would have advantages as efficiency, rapid adaptive reconfiguration and resilience which are essential features for plasticity and cognition (Baronchelli et al. 2013) as well as neurological diseases as stroke and dementia. These advantages would be possible because a failure in a network with such an architecture would most likely affect a peripheral node rather than a hub, as a consequence neither the mean shortest path is strongly increased nor the clustering coefficient decreases. In short, in case of a random failure in the system, e.g. a removal of a node or a malfunctioning given an external or internal perturbation, the information can still reach its final destination in a fast way through other paths. A potential weakness of small-world scale-free networks is the sensitivity in case of targeted attacks, e.g. a hub in the network.

## 5. Conclusion

We proposed a new method to construct brain networks based on k-nearest neighbors (kNN). We showed that kNN-built networks presented small-world and scale-free properties, while the most frequently used method based on density thresholding, did not show them to a significant level: decreased small-worldness in all threshold range as compared to kNN and no scale-free. Considering the small-worldness and scale-free experiments in a large cohort with healthy subjects, *k*∼18 is advisable to construct networks with maximal efficiency.

## 6. Acknowledgements

The authors thank Ulrike von Luxburg and Debarghya Goshdastidar for the discussions and comments.

